# Spatially explicit forest mortality forecasts are driven by autocorrelation, not ecological context

**DOI:** 10.1101/2025.11.19.689366

**Authors:** Keenan Ganz, Liz van Wagtendonk, Pratima Khatri-Chhetri, Monika Moskal

## Abstract

A warmer and drier climate has increased the severity and frequency of drought and insect-induced forest mortality. Forest mortality is an autocorrelated phenomenon that results from a complex interplay between stressors and forest traits. Remote sensing enables us to measure forest mortality in different ways, including aerial surveys; changes in satellite imagery; and individual dead tree mapping in high-resolution imagery. We evaluated the role of autocorrelation and mortality data sources in forecasting drought and insect-induced forest mortality in the western United States. To achieve these objectives, we compared the performance of gradient-boosted regression models against naive autocorrelation models at continental and local scale. Further, we also evaluated bias and variability among observers in aerial surveys. At continental scale, we compared models trained on aerial surveys with a satellite-derived forest mortality dataset. At local scale, we compared models trained on aerial surveys with models trained on maps of individual dead trees. We found that gradient-boosted regression models slightly outperformed naive models across both scales. We also found that covariates related to autocorrelation were the primary drivers of mortality predictions and that model performance declined when they were excluded. This result was consistent across both scales of analysis and all mortality data sources. Furthermore, we found bias in aerial surveys where sites with prior-year mortality were re-surveyed more often than locations without mortality. Our findings showed that forest mortality forecasts based on remote sensing data are not sufficiently accurate to support forest management. To map future mortality risk, we advocate for scaling plot-level mortality models and incorporating accuracy assessments into aerial survey datasets.

## 1 Introduction

Climate change is increasing the frequency and severity of drought and insect-induced forest mortality, as demonstrated by recent compilations of literature and crowdsourced data (International Tree Mortality Network, 2025; Mosig et al., 2024). Mapping tree mortality serves key forest management objectives, including monitoring the forest carbon sink (Migliavacca et al., 2025) and mitigating deleterious wildfire (Stephens et al., 2018). These needs have motivated researchers to centralize mortality observations in global databases, whether derived from remote sensing (Mosig et al., 2024) or field observations (Hammond et al., 2022).

Beyond mapping, forecasting forest mortality could serve two management needs: operational intelligence and driver attribution. Operational intelligence is foresight of imminent forest mortality to direct management activities like thinning and wildfire risk mitigation. Driver attribution is a process to identify the forcings and forest characteristics that were associated with past mortality events to better anticipate future mortality. A common approach to driver attribution is to apply model explanation tools, such as partial dependence plots, to a trained forecasting model (e.g. Preisler et al., 2017). Operational intelligence is a prerequisite for driver attribution; model explanations are only useful when the underlying model has strong predictive power.

Forecast models must be sufficiently accurate to support the above management objectives, but common model performance metrics in scientific literature are not designed around this requirement. For example, regression models that explain less than 70% of the variation in a response variable are equivalent in operational utility (Prairie, 1996). Forest mortality forecasts do not yet meet this benchmark, in part because forest mortality is a multi-year process where external stressors, site characteristics, and tree hydraulic traits interact (Trugman et al., 2021).

Another source of difficulty in forecasting forest mortality is differences among mortality observations. Past work on forest mortality forecasting has relied on aerial surveys (Francis et al., 2025; Preisler et al., 2017) and field observations (McNellis et al., 2021; Rogers et al., 2018) from government monitoring programs. Aerial surveys are performed by human observers, which subjects this data source to inconsistent survey effort, variation among observers, and observer error. Validation efforts found that aerial observers accurately identify the afflicted tree species and mortality-causing agents, however, polygon placement varied considerably among observers (Coleman et al., 2018; Johnson and Ross, 2008; McConnell et al., 2000). Field observations in permanent monitoring plots avoid much of the observation error in aerial surveys but scaling these observations with remote sensing data is complicated by privacy laws that keep forest plot locations confidential.

More recently, individual tree mapping (ITM) has emerged as another source of forest mortality data. ITM uses high-resolution aerial imagery to delineate individual trees and classify each tree as alive or dead (Hemming-Schroeder et al., 2023; Khatri-Chhetri et al., 2024). This approach results in a consistent survey effort over the spatial domain and removes variance among human observers. However, ITM requires sub-meter imagery and considerable computational resources, which precludes widespread adoption of this technique.

In this study, we investigated the role of mortality data sources and spatiotemporal autocorrelation in forecasting drought and insect-induced forest mortality in forests of the western United States. To achieve this objective, we conducted a multi-scale analysis at local and continental scale where we trained gradient-boosted regression models to forecast forest mortality (Table 1; Figure 1) and then assessed how much autocorrelation contributed to forecast skill. At continental scale, we compared aerial surveys with a satellite-derived forest mortality product, while at local scale, we compared aerial surveys with a multitemporal ITM dataset. In both analyses, we compared forecasts against naive models based on autocorrelation. Finally, given the popularity of aerial surveys in measuring forest mortality, we also quantified spatial bias in survey effort and year-to-year spatial variability in mortality polygons.

**Figure 1:**
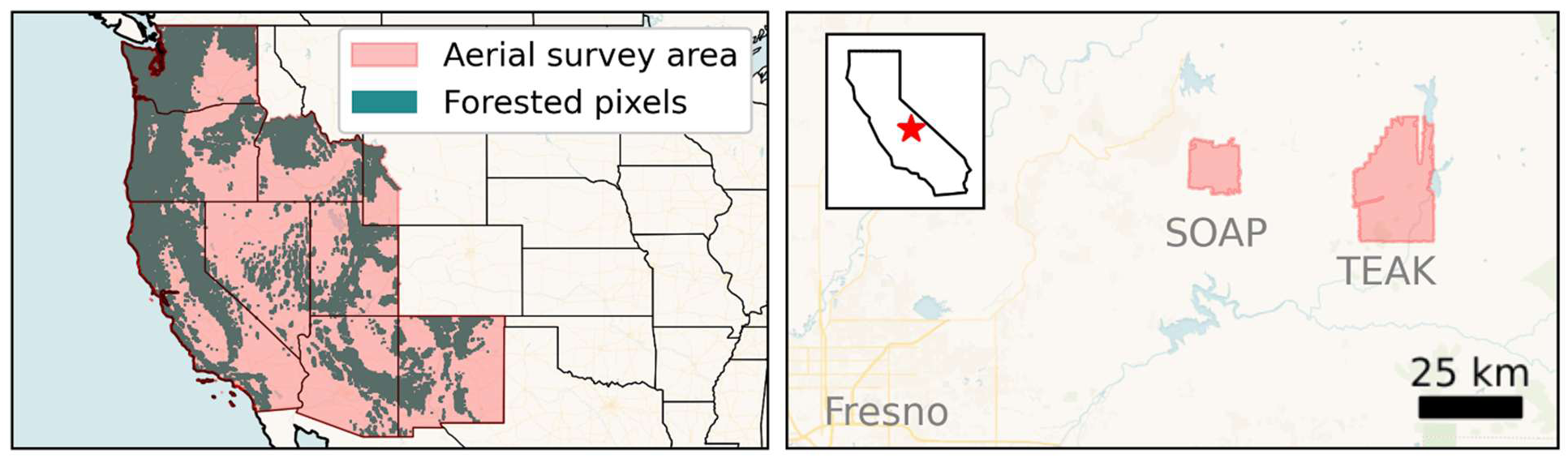
Study areas for our continental-scale (left) and local-scale (right) forest mortality datasets. In the continental-scale dataset, the red region defines the full extent of aerial surveys, but only green forested pixels were considered. SOAP: Soaproot Saddle; TEAK: Lower Teakettle. Basemap tiles were provided by CARTO.

**Table 1:**
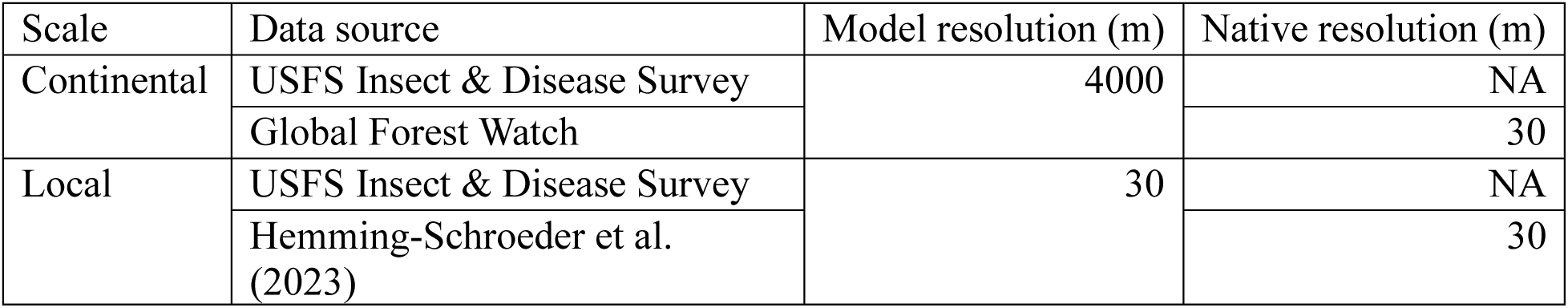
Spatial resolution of forest mortality observations in our local and continental-scale datasets.

| Scale | Data source | Model resolution (m) | Native resolution (m) |
| --- | --- | --- | --- |
| Continental | USFS Insect & Disease Survey | 4000 | NA |
|  | Global Forest Watch |  | 30 |
| Local | USFS Insect & Disease Survey | 30 | NA |
|  | Hemming-Schroeder et al. (2023) |  | 30 |

## 2 Methods

### 2.1 Continental-scale forest mortality data

We developed model-ready forest mortality data through the command-line interface to the Geospatial Data Abstraction Library (Rouault et al., 2025). To ensure this workflow is reproducible and extensible, we managed individual analysis steps with the Python-based workflow engine snakemake (Köster and Rahmann, 2012). Code and model-ready datasets are archived on Zenodo.

We rasterized forest mortality polygons from the US Forest Service (USFS) Insect & Disease Survey in USFS Regions 3, 4, 5, and 6 from 2001 – 2023, including a correction to harmonize different ways surveyors have encoded mortality severity over time (Hicke et al., 2020). We selected mortality polygons attributed to drought, bark beetles, or wood-boring insects; calculated the percent cover of mortality polygons in 4 km pixels, weighted by mortality severity; and masked pixels with less than 10% forest cover. This process resulted in annual raster images from 2001 – 2023 where pixels encode the severity-weighted fraction of forest canopy with mortality (hereafter mortality fraction), and no data if the pixel was unsurveyed or unforested. Documentation for this survey program states that mortality polygons are intended to represent “trees that have died since the last survey” (p.60, McConnell et al., 2000). We therefore interpreted polygons as new mortality, not as an annual snapshot of standing dead trees.

We compared continental-scale forecasts based on aerial surveys with forecasts based on a forest mortality dataset from Global Forest Watch (GFW; Hansen et al., 2013). GFW encodes the year in which a 30 m pixel became unforested between 2000 and 2022 based on satellite imagery. GFW does not rely on human observers and exhaustively measures all forests in our study area, making it an ideal dataset with which to evaluate the role of human observers in mortality forecasting.

We could not limit GFW exclusively to drought and insect-induced forest mortality because this dataset did not attribute forest loss to a damage-causing agent. We reduced the influence of wildfire-induced forest mortality by masking pixels where wildfire occurred within two years of the loss year (Huang et al., 2025) but could not remove the influence of anthropogenic forest loss.

For parity with our aerial survey dataset, we constructed annual loss images for each year of analysis in GFW by selecting 30 m loss pixels for a given year and then regridding to 4 km resolution with mean resampling. This procedure produced imagery of annual new mortality fraction on the same grid as our aerial survey dataset.

### 2.2 Local-scale forest mortality data

In our local-scale models, we compared aerial surveys with an ITM dataset in Hemming-Schroeder et al. (2023) that encoded mortality fraction at 30 m resolution. This dataset is derived from aerial imagery in the Soaproot Saddle and Lower Teakettle sites in the National Ecological Observatory Network. Hemming-Schroder et al. (2023) applied ITM to imagery in 2013, 2017, 2018, 2019, and 2021. To our knowledge, this is the only ITM dataset available that repeatedly measured mortality over a multi-year drought-induced mortality event with minimal influence by wildfire. To facilitate comparison with aerial survey forecasts, we reran our aerial survey rasterization procedure at 30 m resolution within the bounds of the ITM dataset.

### 2.3 Spatial bias in aerial surveys

Aerial surveys varied in their spatial extent. We hypothesized that pixels with mortality are more likely to be surveyed in subsequent years than pixels without mortality. This pattern might introduce bias into the survey data by omitting the beginning of a mortality event following several years without mortality. To test this hypothesis, we calculated the probability ratio

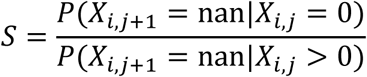

where 𝑋_i,j_ denotes mortality fraction at location 𝑖 and time 𝑗. In a spatially unbiased survey, the survey extent is independent of antecedent mortality observations and 𝑆 = 1, but if a pixel is less likely to be surveyed if mortality was not previously observed there, then 𝑆 > 1. We pooled pixels from all survey years, calculated 𝑆, and tested whether this value significantly deviated from 1 by recalculating 𝑆 on shuffled versions of the original dataset.

### 2.4 Spatial consistency in aerial surveys

Our rasterization procedure obfuscated the spatial pattern of mortality polygons within pixels. Prior work has demonstrated that insect outbreaks spread at a finer scale than our 4 km pixel size (Andrus et al., 2025; Stephenson et al., 2019). Therefore, we hypothesized that autocorrelation in our dataset could arise either from a mortality agent expanding through a pixel or from distinct mortality events in the same pixel (Figure 2). Although the Insect & Disease Survey reports new mortality, polygons may still overlap if a mortality agent partially damages a stand over multiple years. If autocorrelation arises from repeat observations of an expanding mortality event, then we would expect high overlap among polygons over consecutive survey years. This distinction is important because an ideal forest mortality forecast would give advance warning of emergent mortality events. For a model to learn to predict emergent mortality, training data must include instances where previously unobserved mortality occurs.

**Figure 2:**
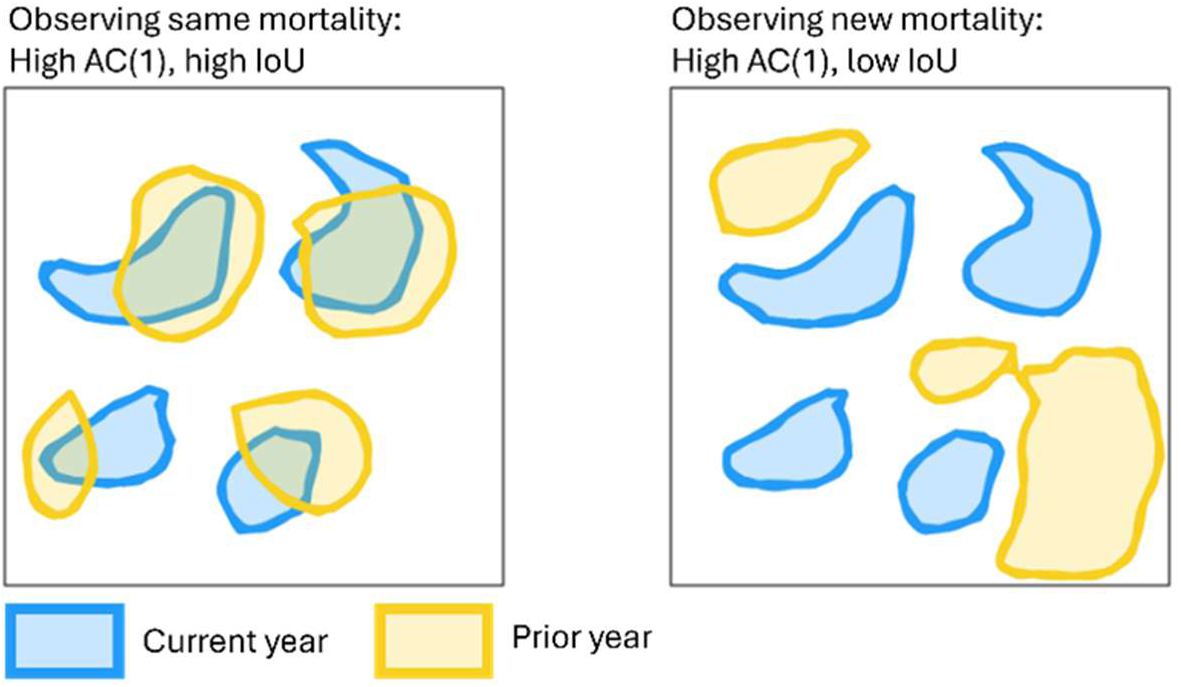
Two hypothesized sources of temporal autocorrelation in aerial surveys. Both panels have similar total mortality area over two consecutive survey years, giving high lag-1 autocorrelation. Overlap among polygons determines intersection over union (IoU), which we hypothesize is higher when repeatedly observing the same mortality event (left panel) versus observing distinct mortality events across survey years (right panel).

In order to assess spatial consistency in aerial surveys, we quantified polygon overlap within pixels as intersection over union (IoU), an area-independent measure of polygon similarity (Figure 2). For each pixel, we calculated IoU for mortality polygons in consecutive survey years attributed to the same host tree species, then calculated mean IoU and lag-1 autocorrelation across all survey years. To reduce the influence of outliers, we only included pixels that were surveyed at least half of the time in this analysis.

Coleman et al. (2018) performed aerial mortality surveys of the same forest at the same time, but with different observers. We used this dataset as a reference point for polygon similarity when forest conditions are constant by calculating the pairwise IoU among all observers (𝑛 = 6 pairs).

### 2.5 Covariate datasets

We trained models to predict future mortality fraction based on annual weather, wildfire area, forest species composition, and topography (Table 2). To quantify annual weather, we calculated seasonal summaries of the monthly Daymet product (Thornton et al., 2022). We calculated summer mean vapor pressure over June – August, minimum winter air temperature over December – February, and water-year precipitation from October to October. Where these periods include two calendar years, the seasonal summary applies to the later year. We quantified wildfire area as the total percent cover of wildfire perimeter polygons in each 4 km pixel. After Preisler et al. (2017), we constructed temporally lagged covariates as listed in Table 2. We assumed that forest species composition and topography were constant through time and simply reprojected these data to our 4 km modeling grid.

**Table 2:** Covariates used to forecast mortality fraction in our continental and local-scale models. Numbers in parentheses indicate the temporal lags, in years relative to mortality, we included in the training data.

| Category | Covariates | Source |
| --- | --- | --- |
| Annual weather | Summer mean vapor pressure (1)<br>Water-year precipitation (1, 2, 3, 4)<br>Minimum winter air temperature (1) | Daymet (Thornton et al., 2022) |
| Wildfire | Total burned area (2, 3, 4) | Monitoring Trends in Burn Severity (Eidenshink et al., 2007) |
| Tree genus basal area | Basal area for *Abies*, *Pinus*, *Populus*, *Tsuga*, and *Pseudotsuga* | Individual Tree Species Parameter Maps (Krist et al., 2014) |
| Topography | Elevation, slope, northness, eastness | ASTER Global DEM (U.S./Japan ASTER Science Team, 2019) |
| Autocorrelation | Prior mortality (1)<br>Maximum prior mortality in adjacent cells (1)<br>Forest area less prior mortality (1) | - |

At local scale we modified our covariate calculation procedure to suit Soaproot Saddle and Lower Teakettle at 30 m resolution. Daymet-derived covariates were too coarse to capture microclimatic variation. We hypothesized that evaporative demand was a function of topography at this scale and represented this source of stress with a heat load index (McCune and Keon, 2002).

Models trained on individual pixels cannot leverage spatiotemporal context to make predictions. Therefore, we also included three autocorrelation covariates that quantified nearby mortality dynamics as mortality fraction in the prior year; maximum mortality fraction among neighboring cells in the prior year; and forest area minus mortality fraction in the prior year (Preisler et al., 2017). This category let us measure the importance of autocorrelation in model performance by comparing the performance of a full model with all categories in Table 2 and a second model with all categories except autocorrelation.

Following initial forecast results, we decided to further investigate the autocorrelation structure of our continental-scale mortality dataset by calculating autocorrelation statistics at a range of spatial and temporal lags. To evaluate temporal autocorrelation, we calculated the Pearson correlation coefficient between pixels at the same location but separated by a temporal lag from 1 to 10 years. To evaluate spatial autocorrelation, we calculated Moran’s 𝐼 for pixel pairs that were from the same survey year and within equally spaced distance bins ranging from 10 km to 60 km. For example, given a distance bin from 10 km to 15 km, we identified all pixel pairs that were separated by between 10 - 15 km and calculated Moran’s 𝐼 from this reduced dataset.

### 2.6 Model training procedure

We trained gradient-boosted regression models to predict mortality fraction with the Python package LightGBM (Ke et al., 2017) in the CryoCloud JupyterHub environment (Snow et al., 2023). Like random forest, gradient-boosted regression is based on an ensemble of decision trees. However, gradient-boosted regression trains decision trees to reduce the current error of the model instead of training on random subsets of the data. This makes gradient-boosted regression more performant than random forest at the cost of a greater risk of overfitting.

We suppose that an operational forest mortality forecast would be run in the same region year after year. In this case our model must generalize across time but not necessarily across space. Therefore, we adopted a temporal cross-validation scheme. At continental scale, we trained models on four sequential years of mortality observations and validated models on the subsequent year. At local scale, we selected three evenly spaced points in 2013, 2017, and 2021. We used 2013 as the baseline for autocorrelation features, 2017 for model training, and 2021 for model validation.

We evaluated model performance according to normalized RMSE and 𝑅^2^ on validation data. We tested for differences in normalized RMSE among models with the Friedman test for repeated samples and post-hoc Wilcoxon signed-rank tests with a Bonferroni correction for multiple comparisons. Since only one validation year was available for our local-scale experiment, we simply report model performance on all validation data.

For our models to be useful, they must outperform naive models based on autocorrelation. At continental scale, we constructed two benchmark naive models: the last mortality observation prior to the validation year and the per-pixel average mortality fraction over the training period. We hereafter refer to these models as last observation and pixel average, respectively. At local scale, the limited temporal extent made the last observation and pixel average models equivalent, so at this scale we only reported results from the last observation model. For the aerial survey dataset at local scale, we calculated the mean mortality fraction per pixel across all surveys prior to 2013 as the pixel average model.

## 3 Results

### 3.1 Spatial bias and variability in aerial surveys

We found that 𝑆 = 1.85, or that pixels without mortality were 85% more likely to be unsurveyed in subsequent years than pixels with nonzero mortality. Permutation testing revealed that this ratio was statistically significant (𝑛 = 499; 𝑝 = 0.002). Per-pixel autocorrelation and mean IoU were positively correlated (𝑟 = 0.37; Figure 3). Surveys of the same forest by multiple observers in Coleman et al. (2018) had geometric mean IoU of 0.33. In our dataset, over 99.9% of all pixels had mean IoU smaller than this value.

**Figure 3:**
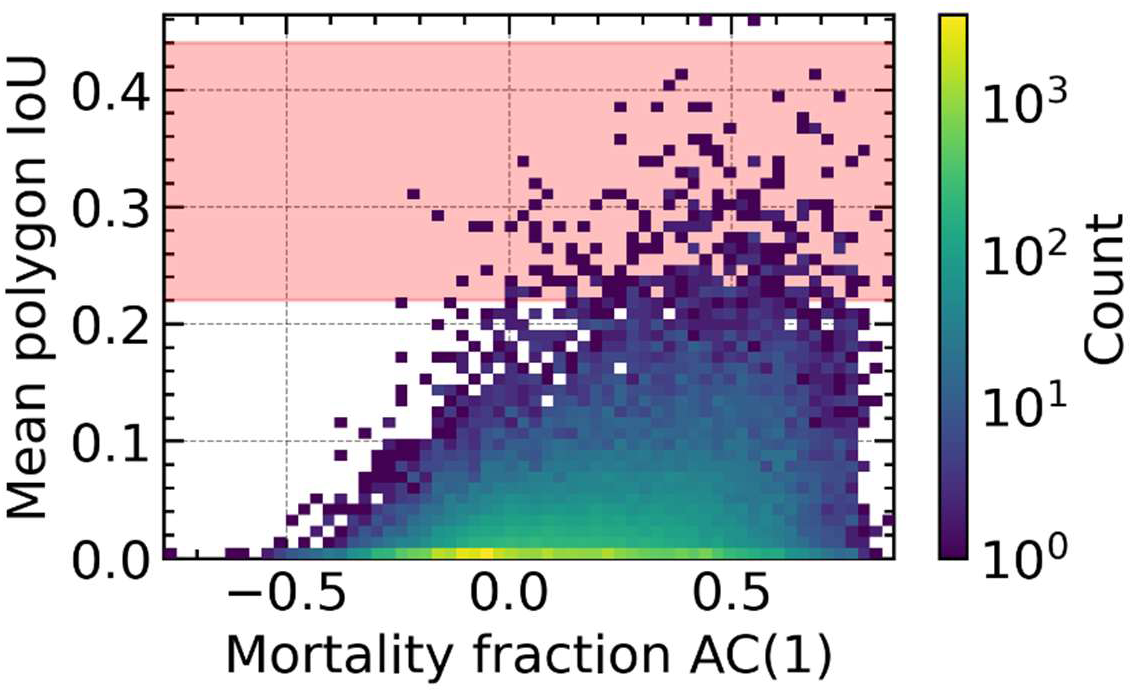
Two-dimensional heatmap of mean polygon intersection over union and lag-1 autocorrelation of mortality fraction. Both metrics were derived from the entire time series for a given pixel. Pixels with fewer than 10 observations over the survey period were excluded. The red region is the range of pairwise IoU among observers from Coleman et al. (2018).

Both continental-scale datasets were autocorrelated in space and time (Figure S1). Spatial autocorrelation on the aerial survey dataset was greater than that for the satellite-derived dataset at all lags, while temporal autocorrelation was comparable between the two datasets.

### 3.2 Continental-scale forecast performance

On the aerial survey dataset, predictive skill differed among models (𝑛 = 19; Friedman’s 𝑄 = 52.6; 𝑝 = 2.19 × 10^-11^), with the full gradient-boosted model performing best (mean NRMSE = 0.92, mean 𝑅^2^ = 0.15). When autocorrelation features were not present during training, the gradient-boosted model performed worse than the full model but slightly better than both naive models (Figure 4). Results were comparable on the satellite-derived dataset, with significant differences among models (𝑛 = 18; Friedman’s 𝑄 = 54.7; 𝑝 = 8.10 × 10^-12^; Figure S2) and the full gradient-boosted model performing best (mean NRMSE = 0.93, mean 𝑅^2^ = 0.13). However, in most validation folds 𝑅^2^ < 0 for both naive models (Figure S3).

**Figure 4:**
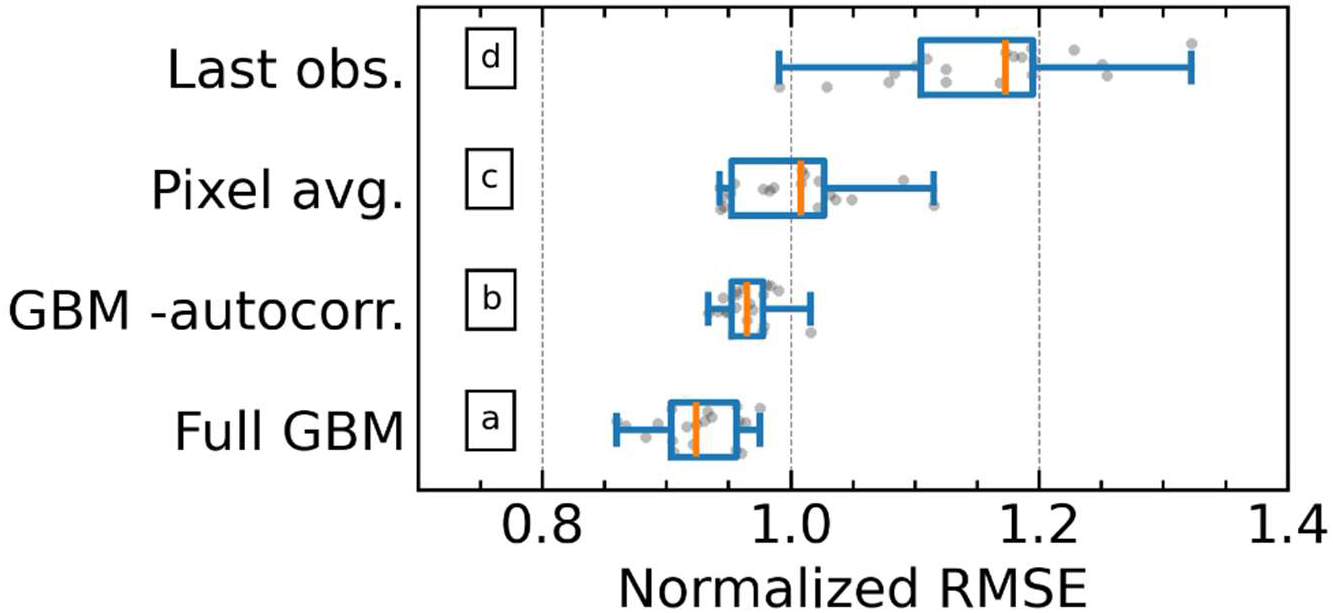
Boxplot of normalized RMSE across validation runs on the continental-scale aerial survey forest mortality dataset. Gray dots show values for individual validation folds with some vertical noise added for legibility. Letter annotations indicate significant differences in performance rank based on Wilcoxon signed-rank tests with Bonferroni correction.

Model performance varied over time on both continental-scale datasets, reflecting the non-stationarity of the prediction problem (Figure S3, Figure S4). Greater variance in model performance on the aerial survey dataset was likely due to differences in survey effort. For example, the surveyed area in 2020 was roughly 70% smaller than the preceding five years and this change coincided with severe degradation in model performance. Although models trained on satellite-derived mortality observations also had degraded performance in 2020, the degradation was not as severe.

Maps of model predictions showed that autocorrelation features were a major driver of model output. Spatial artifacts of the aerial survey process, such as stripes of mortality along the survey flight path, propagated to model output when autocorrelation features were present (Figure 5). However, when autocorrelation features were absent during training, these artifacts disappeared.

**Figure 5:**
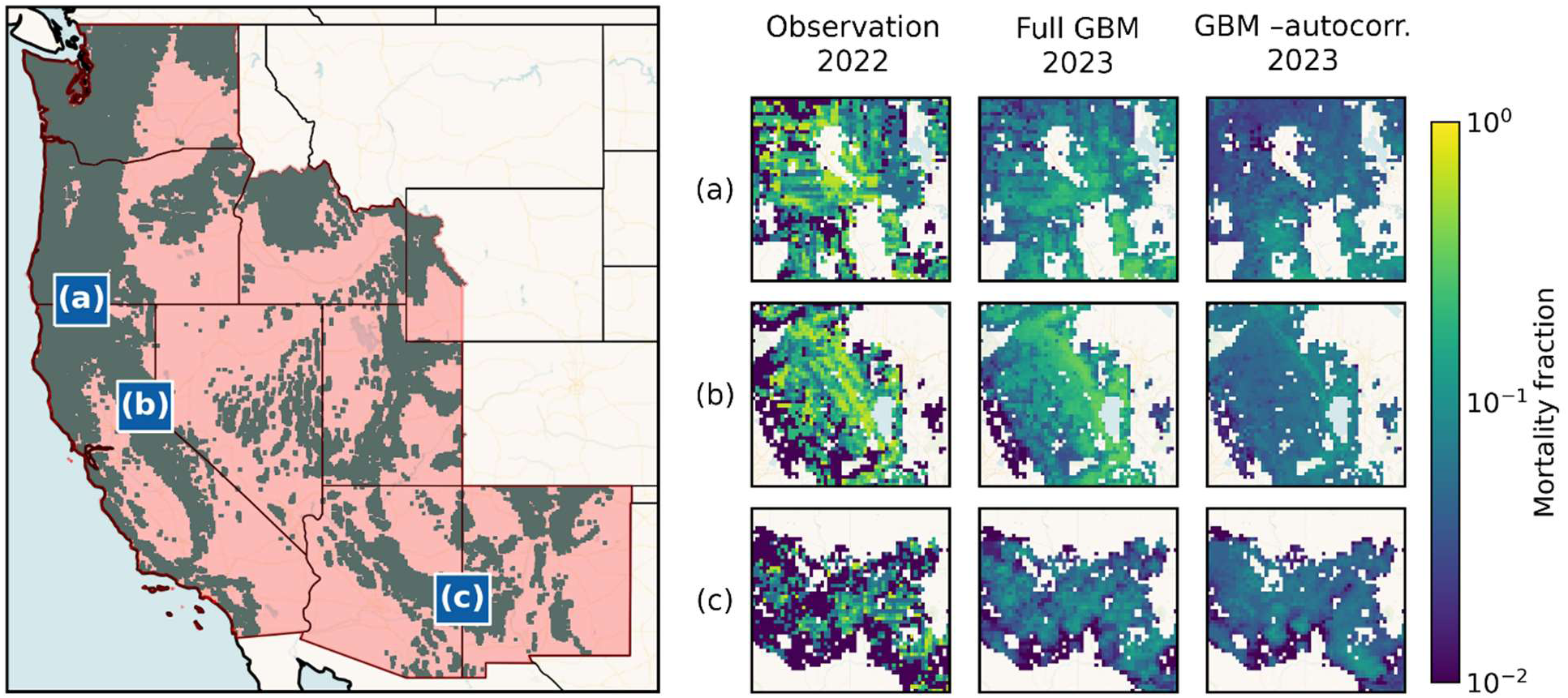
Observed and predicted mortality fraction from the aerial survey dataset at three representative locations in the modeling domain. Blue insets in the left panel show the location of maps in rows (a) – (c) in the right grid. Column labels denote the source of mortality fraction plotted in the right grid. Basemap tiles were provided by CARTO.

### 3.3 Local-scale forecast performance

Forecasts derived from aerial surveys were much less performant at local scale than at continental scale (Table 3). All aerial survey models had 𝑅^2^ < 0. Of these, the long-term pixel average performed best by approximating the overall mean of the data. In comparison, on the ITM dataset the full gradient-boosted model slightly outperformed the naive last observation model, but the gradient-boosted model without autocorrelation features did not.

**Table 3:** Validation metrics for local-scale forecast models trained on mortality data in Soaproot Saddle and Lower Teakettle. NRMSE: normalized root mean square error. Bolding indicates the best-performing model on each dataset.

|  | USFS Insect & Disease Survey |  | Hemming-Schroeder et al. (2023) |  |
| --- | --- | --- | --- | --- |
| Model | NRMSE | $R^2$ | NRMSE | $R^2$ |
| Full GBM | 4.09 | -15.8 | **0.72** | **0.48** |
| GBM -autocorr. | 3.88 | -14.0 | 0.93 | 0.14 |
| Last observation | 4.86 | -22.6 | 0.76 | 0.42 |
| Pixel average | **1.02** | **-0.05** | - | - |

## 4 Discussion

Our results demonstrated that forest mortality forecasts are driven by autocorrelation at both continental and local scale. As measured by 𝑅^2^, model performance degraded by more than half when autocorrelation features were excluded. Autocorrelation is not necessarily undesirable. Indeed, spatiotemporal autocorrelation is a useful property in many prediction problems. However, this result indicates that mortality forecasting models do not support the management objectives laid out in the introduction. Our continental-scale models predict high mortality risk where mortality has already occurred (Figure 5), and no model here meets the benchmark for operational utility from Prairie (1996): 𝑅^2^ > 0.70. Managers would gain similar insights by studying previous mortality surveys. In that sense, our models do not add operational intelligence Moreover, since our models were driven by the spatiotemporal context, they contribute little to our understanding of how other forest conditions and environmental forcing drive mortality dynamics.

We found that spatial and temporal autocorrelation were of similar magnitude on both continental-scale datasets (Figure S1). However, spatial autocorrelation was greater in aerial surveys than in satellite observations of mortality. We suspect that the greater prevalence of autocorrelation in aerial surveys resulted in greater model performance in some validation folds (Figure S3, Figure S4).

The nature of aerial surveys is likely contributing to some of our models’ dependence on spatiotemporal autocorrelation. Virtually none of the pixels in our dataset had IoU consistent with constant forest conditions. However, polygon overlaps increased with temporal autocorrelation. Note that our analysis of IoU only counted overlaps between polygons with the same host tree species. If aerial observers called the host species accurately in mixed forests, autocorrelation in our dataset did not arise from successive mortality events that affected different hosts. Survey effort varied from year to year, and our analysis found that surveyors were less likely to revisit regions without mortality than regions with mortality. This pattern might reflect an operational need to monitor whether past mortality events have spread, but would tend to omit emergent mortality events from being surveyed. Therefore, our mortality dataset may not include the beginning of multi-year mortality events, which would impede our ability to forecast forest mortality where it has not already occurred.

Our analysis of a satellite-derived forest mortality dataset let us examine model performance when autocorrelation was reduced. Global Forest Watch exhaustively observes all pixels in our modeling domain each year and prevents double-counting of mortality events in consecutive survey years. We suspected these factors would reduce autocorrelation in this dataset, and indeed, most naive models had 𝑅^2^ < 0. In this case a model based on biophysical covariates outperformed both naive benchmarks but had only half the predictive power of a model with autocorrelation covariates included (Figure S3). In contrast, the biophysical model did not outperform either naive benchmark on the aerial survey dataset. Although autocorrelation is a significant driver of model output, its importance diminishes depending on how forest mortality data are collected.

Forecast performance also depended on the scale of analysis. Our results indicated that aerial surveys were inappropriate for forecasting forest mortality at local scale (Table 3). Johnson and Ross (2008) found “moderate agreement” between aerial surveys and ground observations at a spatial tolerance of 50 m and 500 m. In our initial 4 km pixel size, mortality polygons could shift by 500 m without affecting adjacent pixels. However, at 30 m resolution, such shifts could drastically affect our models’ ability to learn mortality dynamics. We found that the best prediction of aerial survey-based forest mortality in 2021 was the per-pixel average of surveys that occurred prior to 2017. This result is inconsistent with Hemming-Schroeder et al. (2023), who documented that over 30% of all living trees in 2013 had died by 2021 in the study area.

ITM data do not resolve the limitations of aerial surveys for local-scale analysis. When we trained models on the ITM dataset, we found similar results as our continental-scale analysis of aerial surveys: the full GBM outperformed the naive baseline, but a purely biophysical model did not. Given the limited spatiotemporal extent of the ITM dataset, this result could be attributed to different sources of mortality between the training period and the validation period. This is certainly true for Soaproot Saddle, where wildfires drove most new mortality from 2017 – 2021, while drought was responsible for most mortality from 2013 – 2017 (Hemming-Schroeder et al., 2023).

Aside from the source of mortality observations, forest mortality models are also limited by inherent difficulty in forecasting ecological phenomena. Bark beetles and other wood-boring insects have distinct life cycles and relationships with host tree species that drive mortality dynamics (Stephenson et al., 2019; Trugman et al., 2021). Mortality products based on remote sensing do not necessarily identify the host and cause of mortality, requiring models to disentangle this complexity. Moreover, ongoing climate change increases the difficulty of ecological forecasting by pushing forest ecosystems outside of their historical range of variability.

Given the ecological complexity of forest mortality, high-fidelity mortality observations are key to improving predictive models. Field observations, although expensive, are subject to fewer sources of error than remote sensing products, and mortality forecasts based on field data are feasible (McNellis et al., 2021; Vázquez-Veloso et al., 2025). However, extrapolating these models is difficult because privacy laws preclude sharing precise field plot locations and key forest demographic parameters are unavailable at continental scale. Remote sensing has a role to play in overcoming the limited extent of field observations to enhance our understanding of forest mortality, but our results demonstrate that forest mortality products based on remote sensing alone are insufficient for this aim. Although sensor-fusion approaches for estimating mortality fraction from satellite imagery show promise (Schiefer et al., 2023), aerial surveys will remain critical for near-term and retrospective forest mortality research.

Given the long history of aerial survey programs, we do not hope to relitigate questions around observer accuracy that have already been addressed in prior research (Coleman et al., 2018; Johnson and Ross, 2008). Instead, we suggest that data curators incorporate accuracy assessment, a ubiquitous component of remote sensing (Congalton, 1991), into aerial survey data products. Documentation for the Insect & Disease Survey suggests internal accuracy assessment steps, such as ground-check flights and disputation of erroneous observations (McConnell et al., 2000). However, whether these efforts occur and their findings are unavailable in public-facing data products. In addition to internal accuracy assessment, we also note that US Forest Service performs aerial surveys in parallel with its forest census program Forest Inventory and Analysis. This presents an opportunity for assessing surveyor accuracy against ground surveys, even if results are aggregated to protect landowner privacy. Accuracy assessments based on existing data collection infrastructure would help modeling studies such as our own account for the influence of human error and derive additional insight from the aerial survey program.

In summary, we showed that forest mortality forecasting models are driven by spatiotemporal autocorrelation in mortality observations at both local and continental scale. Although biophysical parameters contributed to model skill, these covariates were less predictive than naive models that were not based on machine learning. We also found evidence that autocorrelation in aerial surveys is in part caused by surveyors’ tendency to reobserve past mortality and place polygons in the same location. However, models remained dependent on autocorrelation when trained on a satellite-derived forest mortality dataset with consistent survey effort and reduced autocorrelation, suggesting that spatiotemporal context is integral to forest mortality dynamics. To improve our understanding of forest mortality, we advocate for incorporating accuracy assessments into aerial survey data products. This could be achieved by publicizing internal accuracy assessment procedures or by coordinating aerial surveys with existing forest census programs.

## Data Availability Statement

Code used to generate the benchmark mortality datasets, figures, and other results in this manuscript is available at [Redacted for anonymity]. A snapshot of code and data products at the time of submission is archived at 10.5281/zenodo.17644170.

## Acknowledgements

This material is based upon work supported by [Redacted for anonymity]. We thank Tom Coleman for providing spatial data to support our analysis of spatial consistency in aerial surveys and the CryoCloud team for providing computational resources for this project.

## Supplementary Information

Calculating the survey probability ratio S.

For two events 𝐴 and 𝐵, the probability of 𝐴 given that 𝐵 is true can be written as

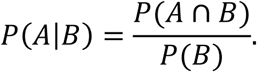

If 𝐴 and 𝐵 are independent, we have 𝑃(𝐴 ∩ 𝐵) = 𝑃(𝐴)𝑃(𝐵) and 𝑃(𝐴|𝐵) = 𝑃(𝐴). Using these definitions, we can rewrite the probability ratio 𝑆 as

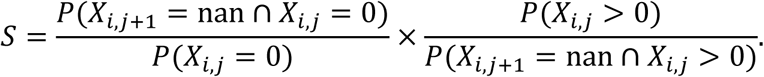

We hypothesized that survey likelihood was dependent on antecedent mortality fraction at that pixel. To test this hypothesis, we identified all pixels in our dataset that were surveyed at least once and constructed boolean masks corresponding to the events in our definition of 𝑆. Note that we discarded the first time step because these data had no antecedent mortality fraction. The masks let us empirically calculate the above probabilities and the value of 𝑆. Then, to test the significance of this value, we repeatedly calculated 𝑆 after shuffling our data in space and time to destroy the observed relationship between survey probability and antecedent mortality fraction.

**Figure S1:**
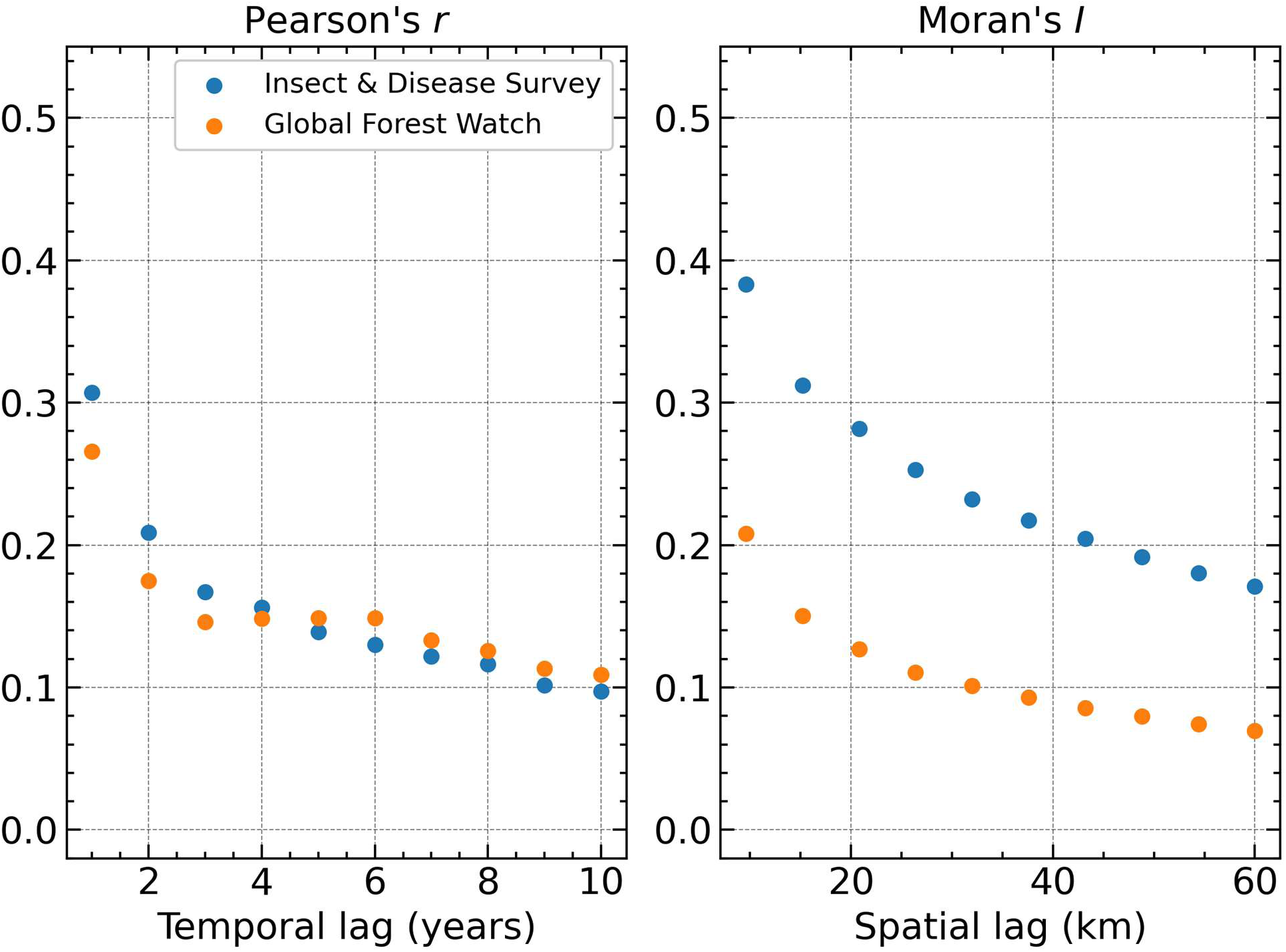
Spatial and temporal autocorrelation statistics for the continental-scale mortality datasets. Lags on the x-axis indicate pixel pairs that were used in the calculation of the relevant statistic. For example, Pearson’s 𝑟 at a temporal lag of two years is the correlation coefficient for all pixel pairs at the same location but separated by two years.

**Figure S2:**
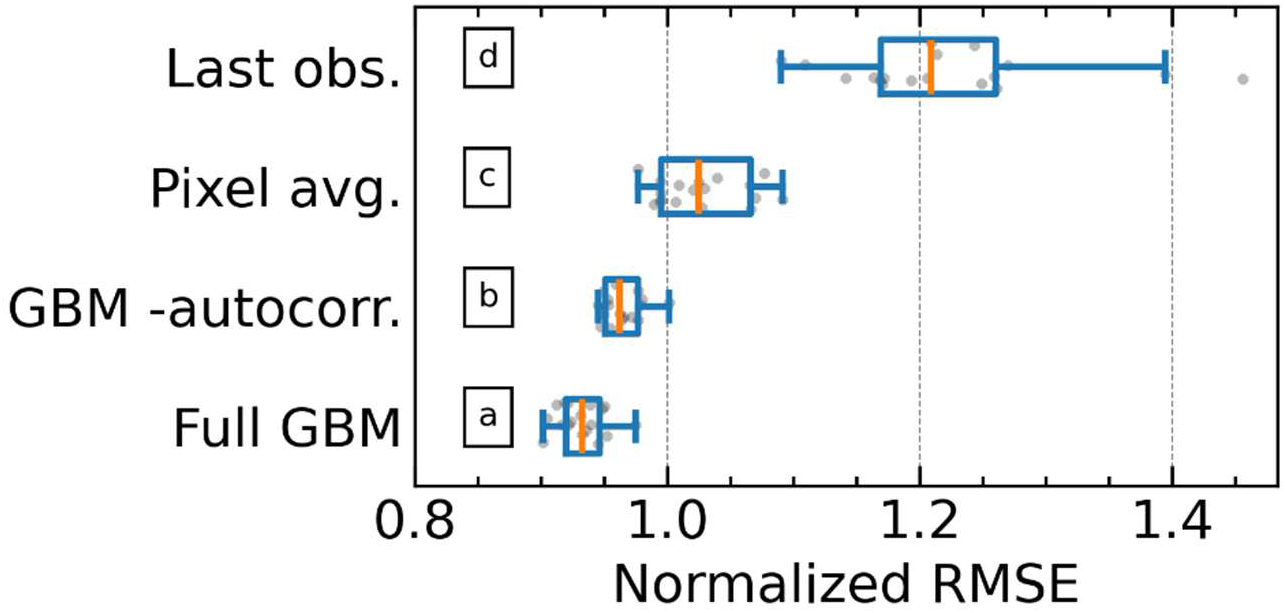
As in Figure 4, but for the satellite-derived forest mortality dataset.

**Figure S3:**
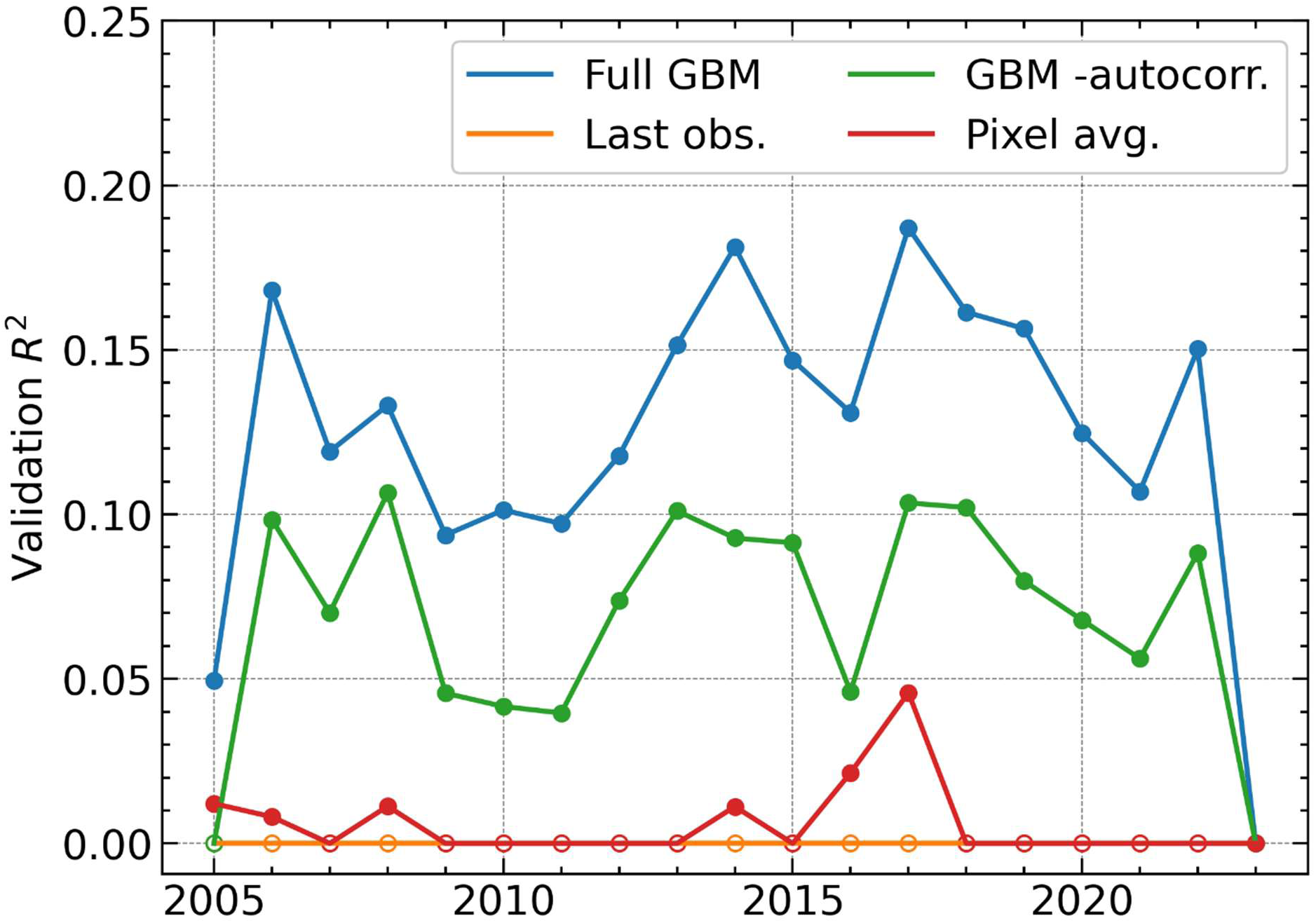
Validation 𝑅^2^ over folds for the continental-scale satellite-derived forest mortality dataset. Points are placed on the validation year. For example, the point at 2010 represents a model trained on data from 2006 – 2009. Open circles along the x-axis denote folds where 𝑅^2^ < 0.

**Figure S4:**
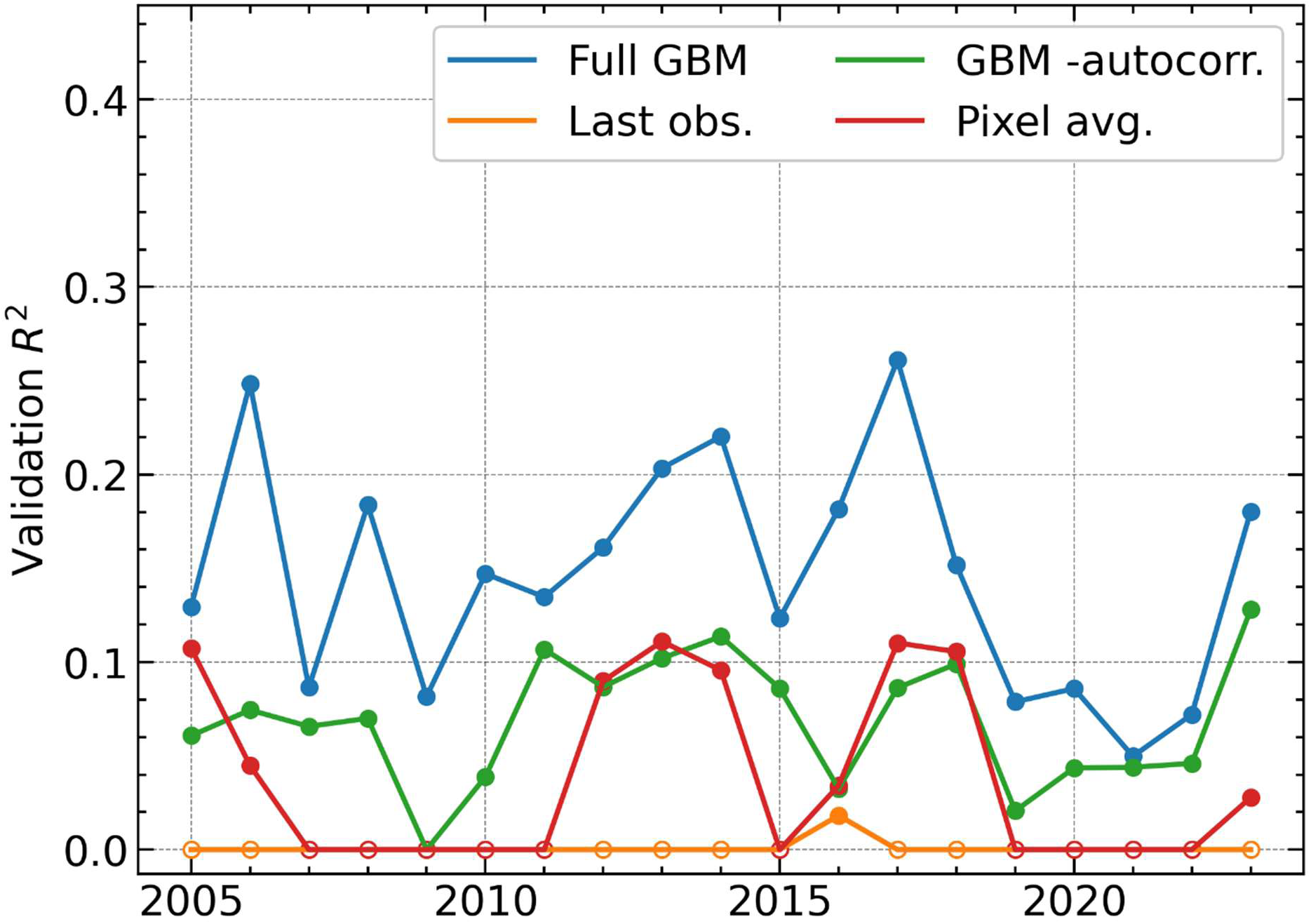
As in Figure S3, except for the aerial survey forest mortality dataset.

